# ProteoformTracker: an interactive tool for planning proteoform detectability in top-down and middle-down proteomics

**DOI:** 10.64898/2026.09.09.750520

**Authors:** Araf Mahmud, Zhihao Zhang, Si Wu, Chen Huang

## Abstract

**Summary:** Characterizing proteome complexity in disease contexts is essential for understanding molecular mechanisms and advancing therapeutic development. Mass spectrometry (MS)-based top-down and middle-down proteomics (TDP/MDP) can resolve intact proteoforms — protein molecules carrying a unique combination of isoform sequence and post-translational modifications (PTMs); however, their technical complexity and modest throughput present challenges for experimental planning and limit their broader application. Here, we present ProteoformTracker, an online web tool that prospectively models MS signal and evaluates the feasibility of using TDP/MDP to distinguish a target proteoform from related isoforms and the background proteome. ProteoformTracker takes as input a gene’s annotated isoforms, a novel long-read/assembled transcript, or an rMATS alternative-splicing event, with or without user-specified PTMs, and predicts each proteoform’s MS1 charge-state envelope and exact isotope pattern, scores per-bond MS2 fragmentation propensity, and searches the full reference human proteome for confounding proteins that could share the target’s intact mass or a charge-state m/z peak. ProteoformTracker also supports middle-down workflows via simulated partial protease digestion. Results are rendered as interactive, zoomable MS1 and MS2 visualizations with live resolvability and fragment-ion statistics, letting users incorporate outside evidence into which confounders they compare against. We envision ProteoformTracker as a useful tool for users to plan TDP/MDP experiments targeting specific proteoforms.

**Availability and implementation:** ProteoformTracker is implemented in R (Shiny) with a Python backend for exact mass and isotope-pattern calculation and is freely available at http://www.proteoformtracker.org together with the documentation and a walkthrough. The source code is available at https://github.com/HuangLabAtUAB/ProteoformTracker under an MIT license.

**Supplementary information:** Supplementary data are available.

## 1 Introduction

Understanding the regulation and complexity of proteins is one of the major research subjects in molecular biology, as proteins are the primary executors of biological processes and are implicated in virtually all disease mechanisms. The diversity of the proteome arises not only from alternative splicing (AS) and alternative open reading frame (ORF) usage (giving rise to protein isoforms), but also from post-translational modifications (PTMs). Accordingly, the term *proteoform* describes a single protein molecule with a unique combination of isoform and PTMs (Smith *et al*. 2013). To date, mass spectrometry (MS)-based proteomics remains the only technique able to systematically profile proteoforms at genome-wide scale. Among MS techniques, bottom-up proteomics (BUP) detects protease-digested (most commonly trypsin) peptides and maps them back to their originating protein sequences. Owing to its throughput and scalability, BUP has been prevalent in large-cohort proteomics studies (Li Y *et al*. 2023). However, BUP has inherent limitations for studying protein diversity: peptides spanning exon–exon junctions are underrepresented (Wang *et al*. 2018), and BUP cannot reconcile which PTM sites co-occur on a single protein molecule and search-based identification of sequence-altering events such as translational errors remains limited by PTM mislocalization in bottom-up data (Mahmud *et al*. 2026).

In comparison, top-down and middle-down proteomics (TDP and MDP) analyze intact proteins or large partial-digest peptides rather than short tryptic peptides, giving direct access to the combinatorial landscape of co-occurring sequence variation and PTMs(Toby, Fornelli, and Kelleher 2016). Although profiling the global proteome has traditionally been more challenging with TDP/MDP than with BUP, improvements in MS instrumentation have significantly increased TDP/MDP throughput, as evidenced by recent large-cohort studies(McCool *et al*. 2022, Melani *et al*. 2022). Nonetheless, further enhancing the utility of TDP and MDP requires not only continued technical improvement, but also more careful, data-driven workflow planning to maximize utility within current technical limitations.

Current TDP/MDP deconvolution engines (e.g., UniDec, FLASHDeconv, TopFD) (Marty *et al*. 2015, Jeong *et al*. 2020, Basharat *et al*. 2023) and database-search tools (e.g., TopPIC, ProSight, MSPathFinder) (LeDuc *et al*. 2004, Kou, Xun, and Liu 2016, Park *et al*. 2017) both operate downstream of MS data acquisition and identify proteoforms from already-recorded spectra. Neither answers the pre-acquisition planning question a researcher designing a TDP/MDP experiment might face — is it worth running this sample at all, given the specific isoforms or PTM states of interest? In the multiomics era, this question becomes even more important, as candidate proteoforms can increasingly be nominated from other data modalities, such as transcript isoforms derived from long-read sequencing. It is important to note that answering that question correctly is harder than a fixed mass-difference rule suggests. The MS resolving power scales with m/z, so the mass difference needed for two proteoforms to appear as separate peaks depends on the pair’s own mass, charge state, and isotope distribution. Moreover, even if the target proteoform can be resolved from relevant proteoforms with AS or PTM events, it can still be confounded by a third, entirely unrelated protein whose intact mass or charge-state peak happens to coincide — a risk that is invisible if comparison is only limited to the relevant proteoforms within the same gene or gene family.

In this study, we developed ProteoformTracker, a user-friendly web tool to address both problems and prospectively assess TDP and MDP feasibility for detecting proteoforms of interest. Any of three input paths — a gene’s annotated Ensembl isoforms, a novel transcript from a FASTA sequence, or an rMATS AS event matched against annotated transcript structure — resolves into a shared internal proteoform representation. From there, ProteoformTracker models each proteoform’s MS1 charge-state envelope and isotope pattern, scores MS2 fragmentation propensity per backbone bond using a formula calibrated against real top-down fragment-ion data, and searches the full reference proteome for confounding proteins in both the mass and m/z domains. The portal generates quantitative, visualized outputs that can be combined with other outside evidence (e.g., RNA-seq data) to help curate exactly which confounders are worth comparing against before committing instrument time.

## 2 Features and implementation

### Proteoform inference and representation

ProteoformTracker takes three types of input to specify the base protein isoform sequences, including gene/isoform selection (Ensembl REST, disk-cached), a pasted/uploaded FASTA sequence (spliced-aligned with minimap2 (Li H 2018) against GRCh38 to identify matching known isoforms, independently translated via candidate ORFs from TransDecoder (Grabherr *et al*. 2011)), or an rMATS event (Shen *et al*. 2014) matched against a precomputed genome-wide exon-structure index to recover full-length transcripts for each arm of the event (Mahmud and Huang 2026). PTMs are specified per isoform through a validated text syntax (<residue position>_<amino acid>_<Unimod name>) drawn from a curated Unimod lookup table to generate the target proteoform and relevant proteoforms to be analyzed (**Supplementary Fig. S1**). See **Supplementary Note 1** for more information.

### MS1: charge-state envelope and isotope pattern

For each proteoform, ProteoformTracker predicts a plausible charge-state envelope: small denatured proteoforms use a basic-residue-count ceiling, while proteoforms above roughly 40 kDa or under native conditions generally use a Rayleigh-limit relationship (Fernandez De La Mora 2000). It then computes the exact isotope pattern from the proteoform’s real elemental composition via pyteomics (Goloborodko *et al*. 2013) and IsoSpecPy (Łacki *et al*. 2017). For each charge state, the predicted isotope spacing is compared against the instrument’s own resolving power, 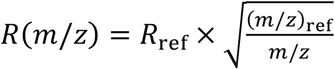, and subsequently the full-width-at-half-maximum peak width is derived as 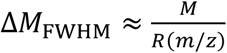 to decide whether the instrument would resolve individual isotope peaks or blur them into one envelope, and the same resolving-power model (with a user-adjustable safety margin, default 1.75 × Δ*M*_*FWHM*_) governs every pairwise peak-collision check in the tool (**Supplementary Figs. S2 and S3**).

### MS2: fragment ladder and fragmentation-propensity scoring

ProteoformTracker generates the full deterministic b/y fragment-ion ladder for every proteoform and scores each backbone bond’s fragmentation propensity with a single logistic regression jointly fit on residue-pair effects (proline/aspartate enhancement), distance to the nearer terminus, local basic-residue (“charge density”), phosphorylation proximity, and proteoform length, calibrated against ≈2.7 million confidently matched b/y-ion observations from two independent public HCD top-down datasets (MassIVE MSV000094311 and MSV000098558) (Forte *et al*. 2024, Sanchez *et al*. 2026). Because a single very long proteoform’s own length can suppress every one of its bonds below fixed tier thresholds, an alternative length-free ranking mode was also provided based on a random forest fit on the same calibration data via ranger (Wright and Ziegler 2017) (**Supplementary Fig. S4B**). See **Supplementary Note 2** for more information.

### Relevant-proteoform comparison

Every proteoform a user checks is compared against every other checked proteoform on two axes: an MS1 charge-envelope overlay (reporting how many charge-state peaks are cleanly separable versus overlapping across the whole set) and an MS2 fragment ladder aligned on a shared exon coordinate axis, with every exon block and every b/y fragment classified as unique, partial, or common depending on how many of the other checked proteoforms share it. A click on any fragment tick computes that ion’s own predicted isotope pattern on demand, overlaid across every other proteoform with a qualifying fragment at the same aligned position (**Supplementary Fig. S4**).

### Confounding-protein search

ProteoformTracker searches a precomputed offline index of the full reference human proteome (UniProt reviewed canonical, n = 20,391 sequences after filtering) for proteins that could confound a chosen target. The search unions two independent axes: a mass-domain window search (candidates whose own intact mass falls within the target’s resolving-power window) and an m/z-domain collision search against a precomputed charge-envelope index (candidates whose charge-state peaks land on the target’s, even when their own intact mass does not overlap with the target’s). Because the m/z-domain search alone can return hundreds of hits, the search results are further prioritized by a lightweight pairwise MS2 overlap count per candidate. Moreover, users can further select which candidates are worth including, using external evidence such as RNA-seq expression, and finalize the confounding-protein list for the full comparison run (**Supplementary Fig. S5**).

### Middle-down support

With middle-down mode selected, ProteoformTracker simulates a limited, partial in-silico digestion (five proteases: OmpT, Lys-C, Lys-N, Glu-C, Asp-N) of every checked proteoform within a user-set mass window, ranks the resulting candidate peptides by feasibility (missed-cleavage count, predicted MS1 peak width) and PTM-site coverage, and applies the identical MS1/MS2 modeling and confounder search — searched against other proteins’ own digest peptides, not intact proteins — to whichever candidates the user selects (**Supplementary Fig. S6**).

### Software

ProteoformTracker is implemented as an R Shiny application (R version 4.5.3), with interactive, client-rendered SVG visualizations (vanilla JavaScript, no charting library dependency) supporting pan/zoom/hover/click across every chart. Exact mass and isotope-pattern calculation is delegated to pyteomics/IsoSpecPy via reticulate. Reference indices (proteome mass index, m/z-collision index, genome-wide exon structure) are precomputed once offline from public UniProt and Ensembl releases and queried at runtime via fast range lookups, not recomputed per request.

## 3 Results and discussion

We have developed ProteoformTracker, a web tool to model MS1 charge envelope and MS2 fragment ladder to assess the feasibility of using TDP and MDP to detect a specific proteoform (i.e., protein isoform + PTM) (**Fig. 1**). The modeling is guided by MS biochemical rules summarized previously (e.g., enhanced cleavage N-terminal to proline and the mobile-proton framework linking basic-residue density to fragmentation propensity) (Wysocki *et al*. 2000, Breci *et al*. 2003) and additionally calibrated against large-scale real MS2 data (**Supplementary Note 2**). The fragmentation-propensity model’s residue-pair, positional, charge-density, length, and phospho-proximity terms were each validated as statistically significant and directionally consistent across two independently acquired public top-down datasets (pooled n ≈ 2.7 million matched b/y-ion observations). This calibration yields tier thresholds with ∼2.5–4.5× fold-enrichment for matched fragment ions over baseline (**Supplementary Fig. S7**). The tool compares a specific target proteoform with two types of proteins separately: relevant proteins that are from the same or similar gene family and share homologous sequence, and confounding proteins that have distinct sequences while sharing similar mass or m/z envelope. This provides the stratified information for identifying the technical caveats in studying a specific AS or PTM from baseline or general protein background. Finally, the tool takes in the input of gene and transcript IDs, FASTA sequence, and AS events and thus can be applied to proteogenomics studies where users have proteoforms derived from genomic and transcriptomic evidence.

**Fig. 1.**
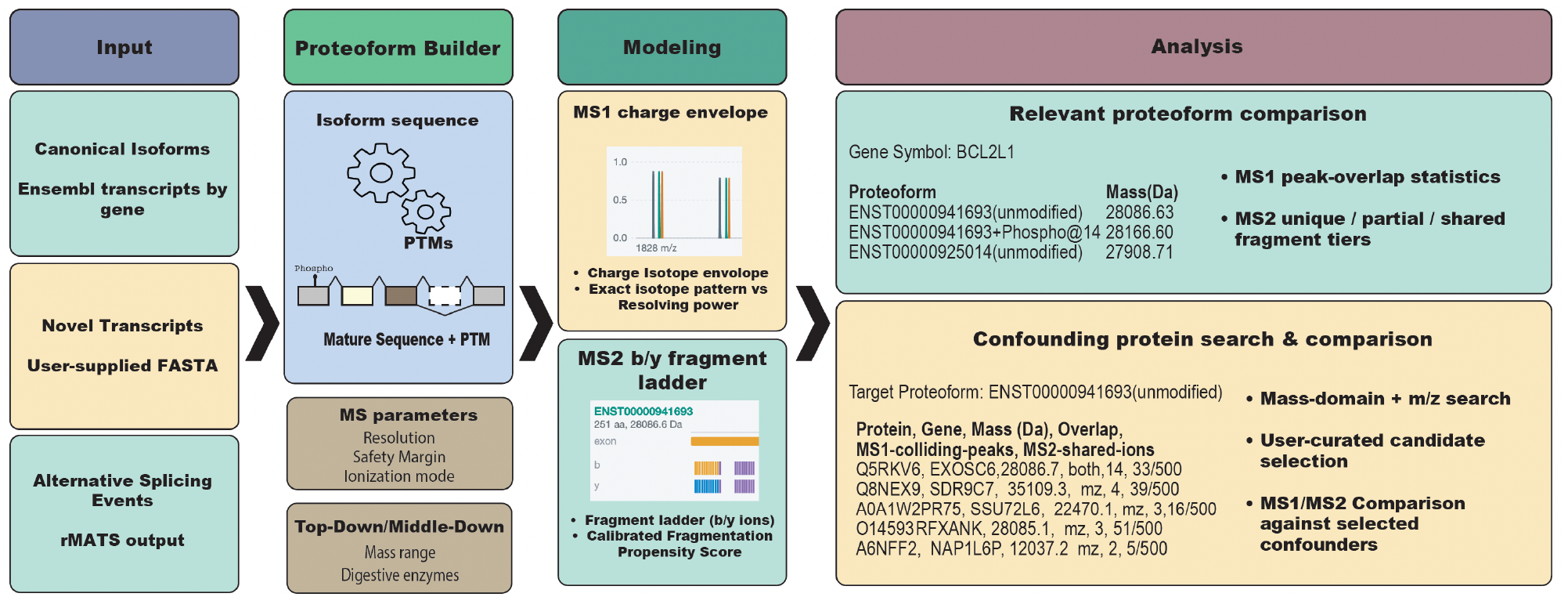
Overview of the ProteoformTracker workflow. First, any of three input paths — a gene’s annotated Ensembl isoforms, a pasted/uploaded FASTA sequence aligned and translated via minimap2/TransDecoder, or an rMATS alternative-splicing event matched against annotated transcripts — resolves into a shared Proteoform object (sequence + PTMs) before downstream scoring runs. Next, each proteoform’s MS1 charge-state envelope and exact isotope pattern are predicted and scored against the Orbitrap resolving-power model; each proteoform’s MS2 b/y fragment ladder is scored for per-bond fragmentation propensity, calibrated against real top-down fragment-ion data. Finally, for the analysis, the relevant protein section compares every checked proteoform against every other (relevant-proteoform comparison); the confounding protein section searches the full reference human proteome for confounding proteins sharing the target’s mass and/or charge-state m/z peaks, lets the user curate which candidates to include using outside evidence, and compares the target against the selected set. Both sections render as interactive, zoomable MS1/MS2 visualizations with live resolvability and fragment-ion statistics.

As a worked example, we applied ProteoformTracker to three BCL2L1 proteoforms — two splice isoforms and a phosphorylated (Ser14) variant of one (**Supplementary Fig. S1B**). At default instrument settings (R = 120,000 @ m/z 200), all three proteoforms are fully resolvable at the MS1 level (**Supplementary Fig. S4A**) and further supported by dozens of proteoform-specific MS2 fragment ions per proteoform (27–31 unique b/y ions each; **Supplementary Fig. S4C**). A confounder search against the full reference human proteome, however, identified 124 unrelated proteins sharing mass and/or m/z with the target isoform; comparing it against the five highest-priority confounders showed that all 17 of its MS1 charge-state peaks collided with at least one confounder (**Supplementary Fig. S5A–B**). Despite this, the target’s MS2 fragment ladder retained 59 unique ions (found in none of the five confounders) and 19 shared with only some of them (**Supplementary Fig. S5C**), indicating fragment-level evidence could still support confident detection when intact-mass evidence is confounded. This illustrates ProteoformTracker’s core utility: quantifying not only whether a target proteoform is distinguishable from its close relatives, but whether that evidence survives a realistic confounding background, and how much supportive evidence can be expected from each acquisition level (i.e., MS1 or MS2).

## Acknowledgements

The authors thank Qi Zhou for constructive discussions in developing this project.

## Data Availability

The fragmentation-propensity model was calibrated against publicly available top-down proteomics datasets deposited in MassIVE under accessions MSV000094311 and MSV000098558. The reference proteome mass index and m/z-collision index are built from the public UniProt reviewed human canonical proteome (UP000005640); the exon-structure index is built from the public Ensembl GRCh38 gene annotation. Scripts to regenerate all reference indices from these public sources are included in the software repository. No new experimental data were generated for this study.

