## Supplementary Figures for "ProteoformTracker: an interactive tool for planning proteoform detectability in top-down and middle-down proteomics"

*All panels in Figs. S1-S6 were captured from a live run of the application (gene BCL2L1, top-down mode) unless noted otherwise. Screen captures illustrate real, computed output, not mockups. Fig. S7 is a rendered statistical plot, not a screenshot.*

### A. Gene/isoform catalog with PTM specification

**Gene symbol**

BCL2L1

Load isoforms

BCL2L1: 40 transcripts in the precomputed exon index, 40 with a real fetched protein sequence, collapsed to 9 unique protein sequence(s).

▼ **Isoform catalog (real Ensembl transcripts for this gene)** (2 of 9 selected)

| | Transcript | Size | Exons | PTMs |
|---|---|---|---|---|
| ☐ | ENST00000450273 | 307 aa, 34112.6 Da | exons 2-4 | e.g. 133_T_Phospho; 210_P_Oxidation; 215_S_Sulfo |

(+1 more transcript(s) with this same sequence: ENST00000941694)

| | Transcript | Size | Exons | PTMs |
|---|---|---|---|---|
| ☑ | ENST00000941693 | 251 aa, 28086.6 Da | exons 2-4 | 14_S_Phospho |
| ☑ | ENST00000925014 | 250 aa, 27908.7 Da | exons 2-4 | e.g. 133_T_Phospho; 210_P_Oxidation; 215_S_Sulfo |

(+1 more transcript(s) with this same sequence: ENST00000925015)

| | Transcript | Size | Exons | PTMs |
|---|---|---|---|---|
| ☐ | ENST00001001206 | 241 aa, 26923.1 Da | exons 2-4 | e.g. 133_T_Phospho; 210_P_Oxidation; 215_S_Sulfo |

(+2 more transcript(s) with this same sequence: ENST00001001209, ENST00001001211)

| | Transcript | Size | Exons | PTMs |
|---|---|---|---|---|
| ☐ | ENST00000307677 | 233 aa, 26032.7 Da | exons 2-3 | e.g. 133_T_Phospho; 210_P_Oxidation; 215_S_Sulfo |

(+16 more transcript(s) with this same sequence: ENST00000376062, ENST00000420488, ENST00000434194, ENST00000439267, ENST00000456404, ENST00000676582, ENST00000676942, ENST00000677194, ENST00000678563, ENST00000870614, ENST00000870619, ENST00000870620, ENST00000870622, ENST00000925016, ENST00000925017, ENST00001133412)

| | Transcript | Size | Exons | PTMs |
|---|---|---|---|---|
| ☐ | ENST00001001212 | 231 aa, 25659.4 Da | exons 2-4 | e.g. 133_T_Phospho; 210_P_Oxidation; 215_S_Sulfo |

### B. Result proteoform table

**Result proteoform table**

Every checked isoform (plus its parsed PTM combinations) is included in the comparison below. Pick one as the confounder-search target:

| Proteoform | Mass (Da) |
|---|---|
| ENST00000941693 (unmodified) | 28086.63 |
| ENST00000941693 + Phospho@14 | 28166.60 |
| ENST00000925014 (unmodified) | 27908.71 |

**Confounder-search target**

ENST00000941693 (unmodified) ▾

Run analysis

Click "Run analysis" to (re)compute MS1/MS2/confounder results for the currently included proteoforms.

### C. FASTA sequence input (Option 2)

#### Input selection

Reset

○ **1. Gene -> isoform -> proteoform** ◉ **2. FASTA sequence** ○ **3. rMATS alternative-splicing results**

#### FASTA sequence input

Provide a single spliced mRNA/cDNA (or CDS) nucleotide sequence -- not raw genomic DNA with introns. Two independent steps run: minimap2 spliced-aligns it against GRCh38 to identify which known gene/isoforms it belongs to (this does not depend on translation at all), and TransDecoder separately finds candidate open reading frames. Pick a known isoform to compare its exon structure against, pick which ORF candidate to use as the translation, add PTMs, then send it into the same comparison view as Option 1.

Paste sequence | Upload file

**Paste FASTA**

>my_transcript
ACGTACGT...

Run sequence analysis

### D. rMATS alternative-splicing input (Option 3)

#### Input selection

Reset

○ **1. Gene -> isoform -> proteoform** ○ **2. FASTA sequence** ◉ **3. rMATS alternative-splicing results**

#### rMATS alternative-splicing results

rMATS only reports the differential exon(s) and their immediate flanking exons, not the rest of the transcript -- so a full-length proteoform can't be computed from the event alone. Instead, ProteoformTracker looks up which already-annotated transcripts of the gene (in the same precomputed exon index Option 1 uses) structurally match each arm of the event (exon-inclusion vs. exon-skipping for SE; 1st-exon vs. 2nd-exon for MXE; intron-retained vs. -spliced for RI; long- vs. short-exon form for A5SS/A3SS), so you get real, full-length proteoforms rather than just the local differential region. Every arm also gets a "constructed" synthetic isoform (a user-pickable backbone transcript with the local region replaced by rMATS' own reported exons), so an arm with no real annotated match still has something usable.

**Supplementary Fig. S1.** Proteoform inference and representation. (A) Gene/isoform selection: real Ensembl transcripts for the queried gene, with a free-text PTM specification box per row. (B) Result proteoform table: every checked isoform, plus one row per valid PTM combination, with its own computed mass; one proteoform is picked as the confounder-search target before running the analysis. (C) Option 2 (FASTA sequence): a pasted/uploaded transcript sequence, independently spliced-aligned (minimap2) and ORF-called (TransDecoder). (D) Option 3 (rMATS alternative-splicing results): an uploaded rMATS event, matched against annotated transcript structure to recover full-length proteoforms for each arm of the event.

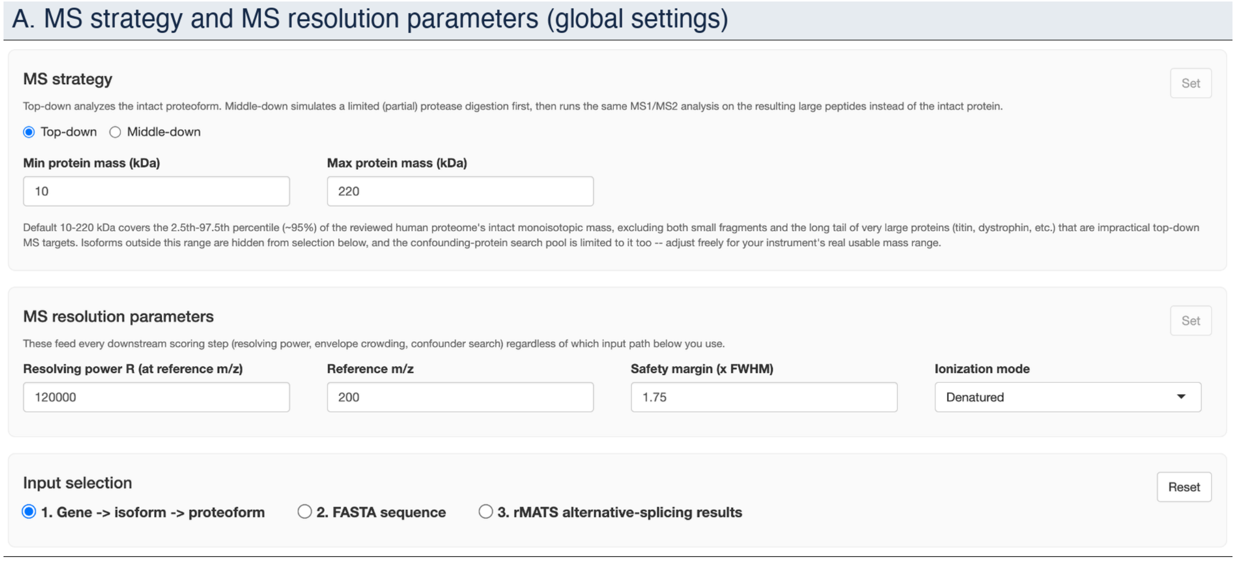


**Supplementary Fig. S2.** Global MS1/MS2 configuration. (A) The MS strategy panel (top-down vs. middle-down, usable mass range) and MS resolution parameters panel (resolving power, reference m/z, safety margin, ionization mode) that parametrize the charge-envelope and resolving-power model described in the main text (see Supplementary Fig. S3 for the model's actual output) — both apply to every input path and must be explicitly confirmed (“Set”) before any analysis can run.

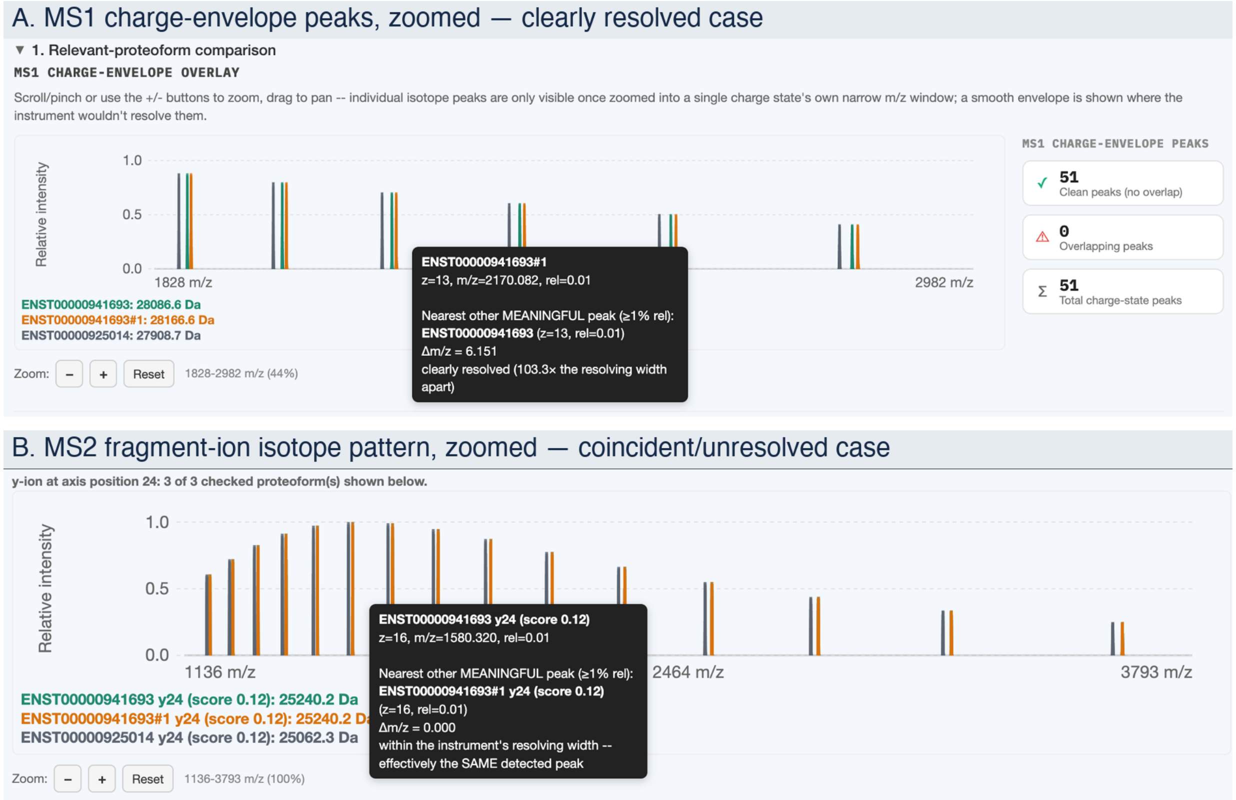


**Supplementary Fig. S3.** Charge-envelope and isotope-pattern modeling: resolved vs. unresolved cases. (A) MS1 charge-envelope peaks, zoomed into a single charge-state cluster; hovering a peak reports the nearest other meaningful peak's Δm/z and whether it clears the instrument's resolving width — here, two proteoforms' peaks sit 103.3× the resolving width apart and are clearly resolved. (B) The same resolvability check applied to one MS2 fragment ion's own predicted isotope pattern (reached by clicking a b/y tick on the fragment ladder, Supplementary Fig. S4C): here two proteoforms' y24 ions coincide exactly (Δm/z = 0.000) and are reported as effectively the same detected peak — the same resolving-power model underlies both the MS1 and MS2 checks.

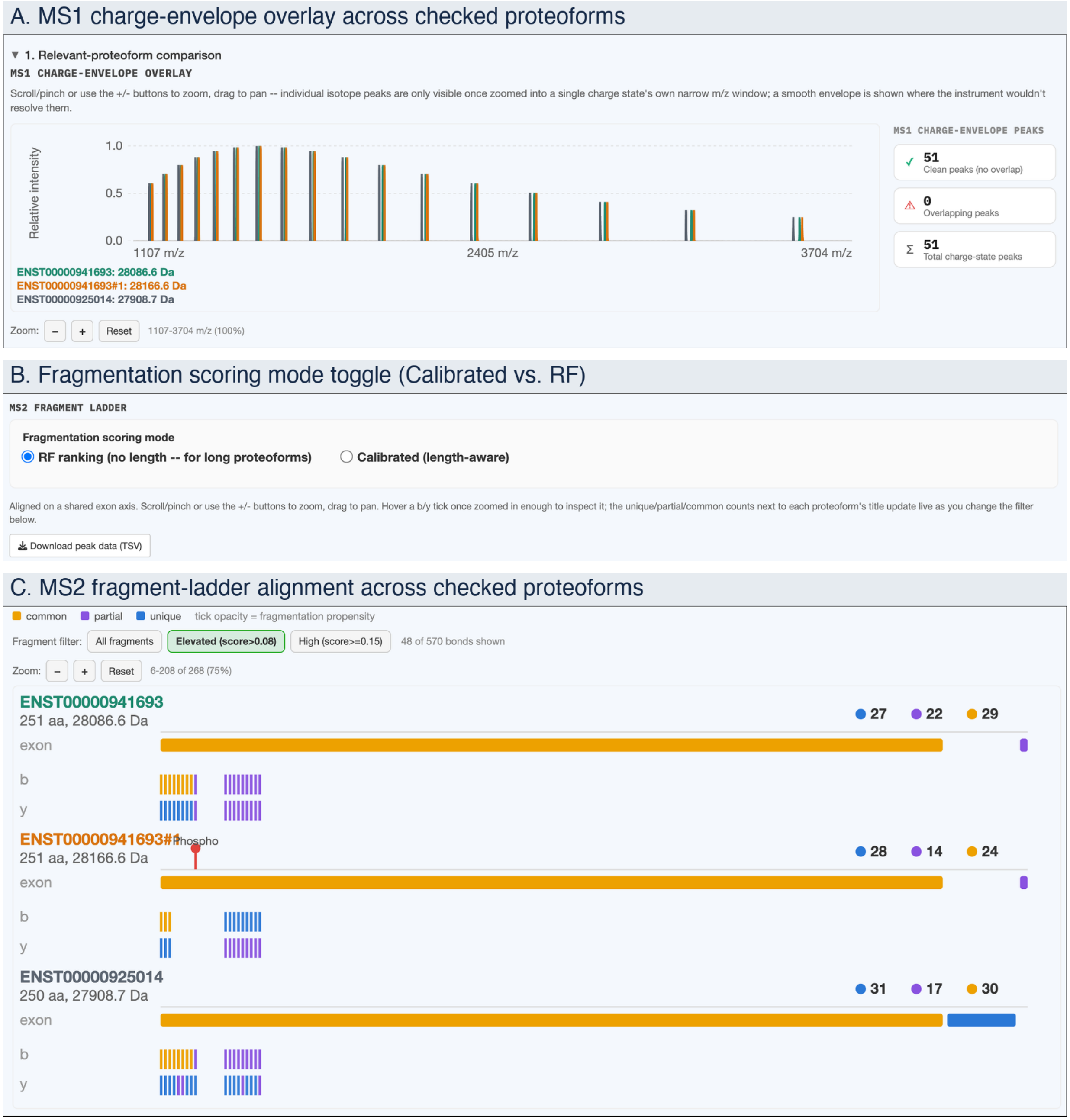


**Supplementary Fig. S4.** Relevant-proteoform comparison. (A) MS1 charge-envelope overlay across every checked proteoform, reporting how many charge-state peaks are cleanly separable versus overlapping. (B) The fragmentation scoring mode toggle: Calibrated (length-aware, the default) vs. RF ranking (length-free, for very long proteoforms whose own length can suppress every bond below the calibrated tier thresholds — see Supplementary Fig. S7 for cross-study validation of both modes), which determines how the ladder in (C) is scored and colored. (C) MS2 fragment ladder for the same proteoforms, aligned on a shared exon coordinate axis; clicking any fragment tick computes that ion's own predicted isotope pattern on demand, overlaid across every other proteoform with a qualifying fragment at the same aligned position (Supplementary Fig. S3B).

### A. Confounding-protein candidate list

▼ **2. Confounding-protein search (single target)**

Target: ENST00000941693#bare. 30 real confounding protein(s) found (0 by mass window, 29 by m/z collision, 1 by both). Showing the **30** highest-priority of **124** total found -- capped to keep the search fast. Select which to include below, then "Compare selected confounders".

Every one of these shares mass and/or m/z with the target -- none are checked by default, so pick the ones worth a full comparison (or use outside evidence, e.g. RNA-seq expression, to guide which to include) before comparing. "MS2 shared ions" is a pairwise count (this candidate alone vs. the target, out of the target's own total b/y ion count) -- the multi-way unique/partial/common tiers shown after comparing are computed fresh from whichever candidates you leave checked, not from this number directly.

Select all | Select none

| | Candidate | Gene | Mass (Da) | Found via | MS1 colliding peaks | MS2 shared ions (vs. target alone) |
|---|---|---|---|---|---|---|
| ☑ | Q5RKV6 | EXOSC6 | 28086.7 | both | 14 | 33 / 500 |
| ☑ | Q8NEX9 | SDR9C7 | 35109.3 | mz | 4 | 39 / 500 |
| ☑ | A0A1W2PR75 | SSU72L6 | 22470.1 | mz | 3 | 16 / 500 |
| ☑ | O14593 | RFXANK | 28085.1 | mz | 3 | 51 / 500 |
| ☑ | A6NFF2 | NAP1L6P | 12037.2 | mz | 2 | 5 / 500 |
| ☐ | O15120 | AGPAT2 | 30894.1 | mz | 2 | 36 / 500 |

Compare selected confounders

### B. MS1 charge-envelope comparison vs. selected confounders

▼ **MS1 charge-envelope overlay**

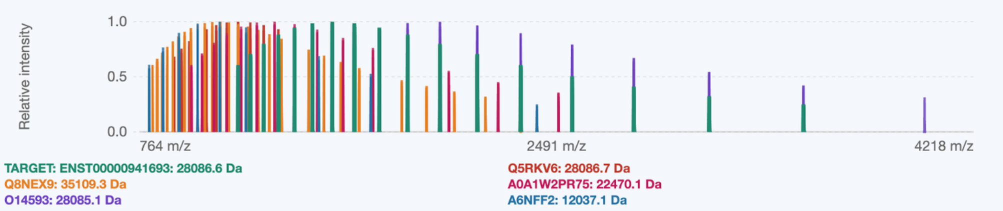


TARGET'S MS1 PEAKS

✓ **0** Clean peaks (no overlap)

⚠ **17** Overlapping peaks

Σ **17** Total charge-state peaks

Zoom: – + Reset 764-4218 m/z (100%)

### C. MS2 fragment ladder vs. selected confounders

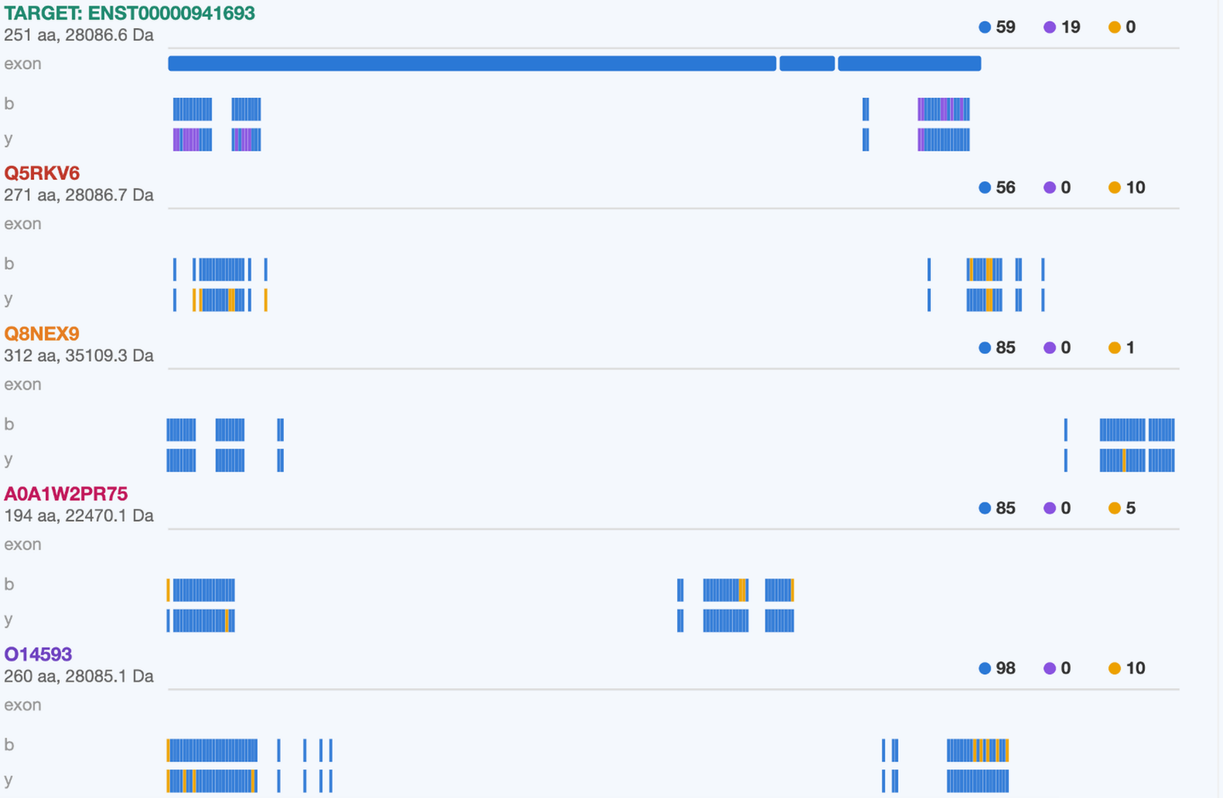

**Supplementary Fig. S5.** Confounding-protein search. (A) Candidate list from the precomputed reference-proteome index: each candidate shows its UniProt accession, gene symbol, mass, how it was found (mass-domain window, m/z-domain collision, or both), and a lightweight pairwise MS2 overlap count; checkboxes let the user curate which candidates proceed to the full comparison using external evidence. (B) MS1 charge-envelope comparison of the target against the selected confounding proteins. (C) MS2 fragment ladder for the target, tiered against the selected confounding proteins.

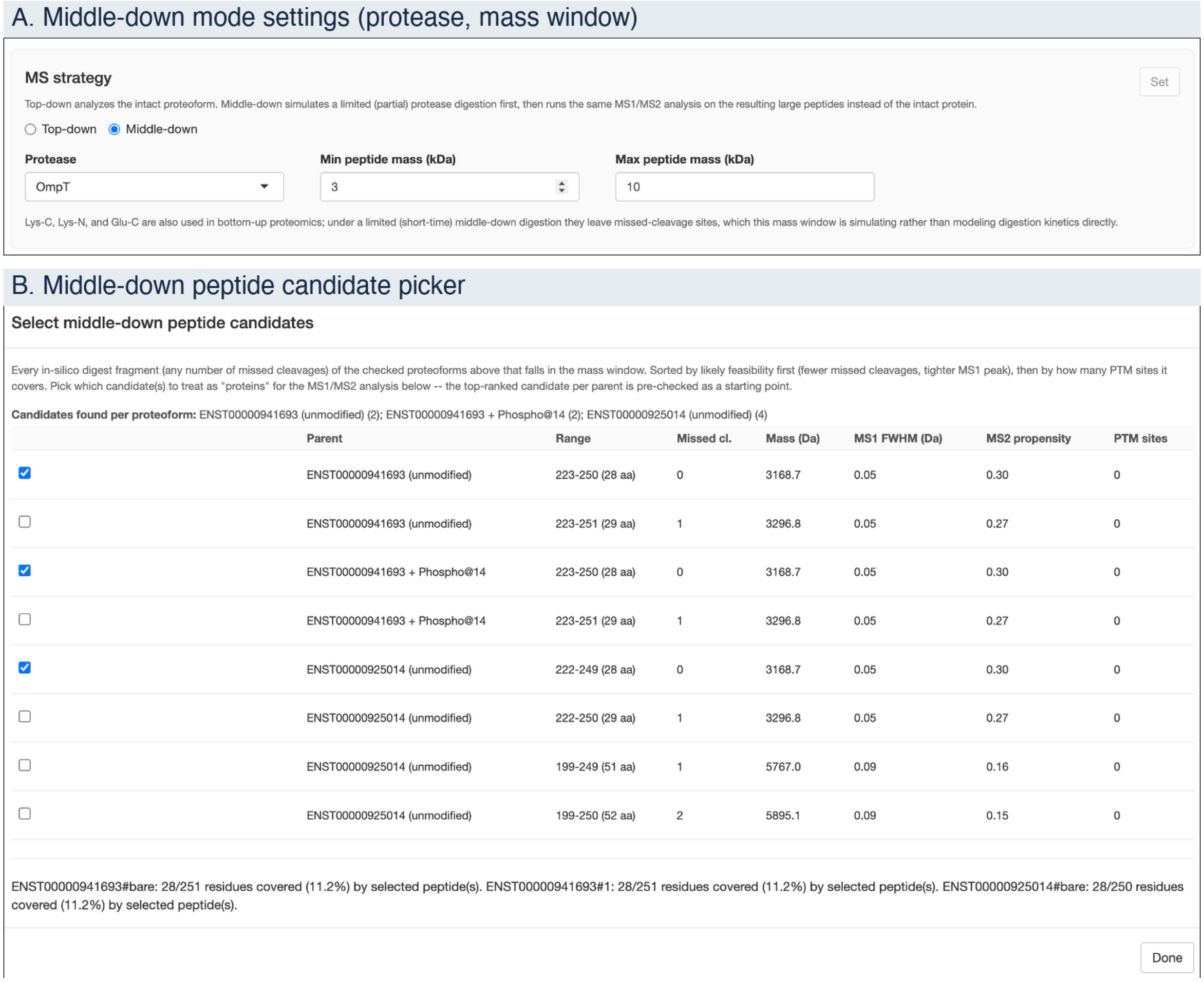


**Supplementary Fig. S6.** Middle-down support. (A) Middle-down mode settings: protease choice (OmpT, Lys-C, Lys-N, Glu-C, or Asp-N) and the peptide mass window used to filter in-silico digest fragments. (B) The resulting peptide candidate picker: every in-silico digest fragment within the current mass window is ranked by feasibility (missed-cleavage count, predicted MS1 peak width) and PTM-site coverage; checked candidates are treated as “proteins” for all downstream MS1/MS2 modeling and confounder search.

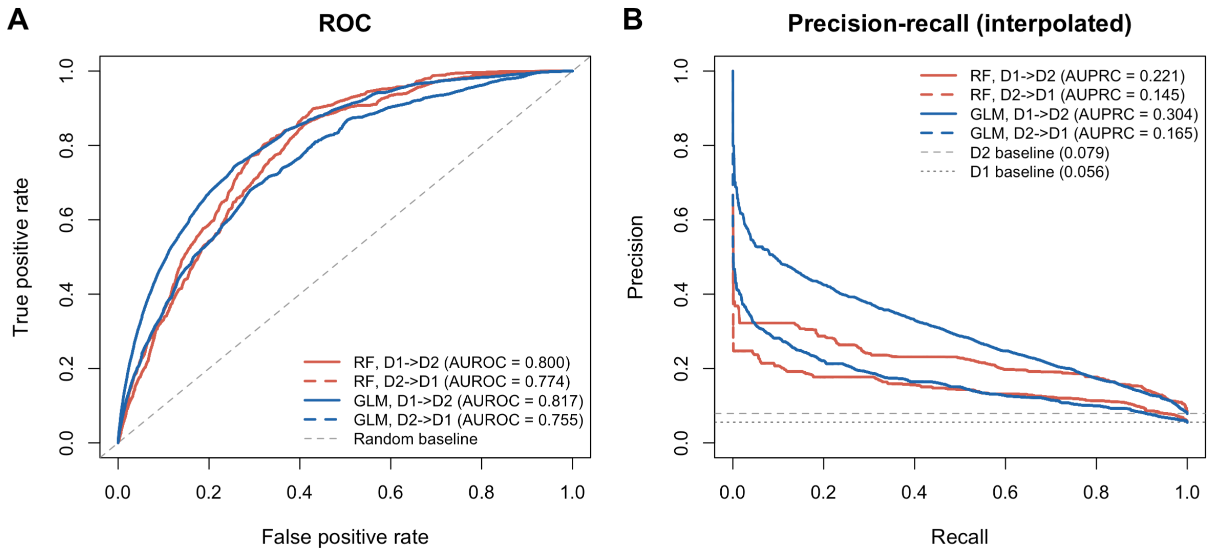


**Supplementary Fig. S7.** Cross-study validation of the fragmentation-propensity scoring models. Discrimination of the calibrated GLM (length-aware) and RF (length-free) fragmentation-propensity scoring modes, each refit from scratch on one MassIVE dataset in its entirety and evaluated on the other, fully unseen dataset, in both directions. (A) Receiver-operating-characteristic (ROC) curves. (B) Precision-recall curves shown as the monotone interpolated envelope (precision at each recall replaced by the maximum precision achievable at that recall or higher; AUPRC values are computed from the raw, non-interpolated curve and are unaffected by this display choice), with dashed/dotted lines marking each held-out test set's own baseline match rate (D2: 7.9%; D1: 5.6%). In both (A) and (B), solid lines are trained on MSV000094311 (D1), evaluated on MSV000098558 (D2); dashed lines are trained on MSV000098558 (D2), evaluated on MSV000094311 (D1). GLM leads RF on both metrics in the D1→D2 direction (AUROC 0.818 vs. 0.800; AUPRC 0.304 vs. 0.221); in the reverse direction RF leads on AUROC (0.774 vs. 0.755) while GLM retains the lead on AUPRC (0.165 vs. 0.145) — neither scoring mode dominates uniformly across both holdout directions, and both substantially exceed the random/baseline reference throughout.
