## Supplementary Note for "ProteoformTracker: an interactive tool for planning proteoform detectability in top-down and middle-down proteomics"

### Supplementary Note 1: Input parsing and proteoform inference

ProteoformTracker resolves each of its three input paths into a common internal Proteoform representation (mature amino-acid sequence plus a list of applied PTMs) before any downstream MS1/MS2 scoring runs, following the same general strategy as IsoPepTracker (Mahmud and Huang 2026) for reconciling heterogeneous transcript-level inputs into concrete protein sequences. An overview of this pipeline is shown in **Supplementary Fig. N1**.

For the **gene → isoform** path, ProteoformTracker queries the Ensembl REST API live for a given gene symbol, retrieving every annotated transcript's coding sequence and translating it to its mature protein sequence; results are cached to disk per transcript so repeated queries for the same gene do not re-hit the network.

For the **novel transcript** path, a user-submitted FASTA sequence (from long-read sequencing or de novo assembly) is processed in two independent steps. First, minimap2(Li 2018), run with splice-aware parameters against the GRCh38 genome, aligns the sequence and identifies which known gene and annotated isoform(s) it structurally resembles. Independently, TransDecoder (Grabherr *et al.* 2011) identifies candidate open reading frames (ORFs) from the raw sequence based on coding potential. The user selects which known isoform to compare exon structure against and which ORF candidate represents the correct translation.

For the **rMATS alternative-splicing event** path, ProteoformTracker takes an rMATS results file, and for each arm of a selected event, matches the event's differential and flanking exon coordinates against a precomputed genome-wide exon-structure index (built once, offline, from an Ensembl GTF via rtracklayer) to identify which already-annotated transcript(s) of the gene structurally correspond to that arm. This recovers real, full-length proteoform sequences rather than only the local differential region rMATS itself reports.

Across all three paths, PTMs are applied on top of the resolved isoform sequence through a validated text syntax (<residue position>_<amino acid>_<Unimod name>), checked against the real residue at that position and resolved against a curated Unimod lookup table, before the final Proteoform object is passed to MS1/MS2 scoring.

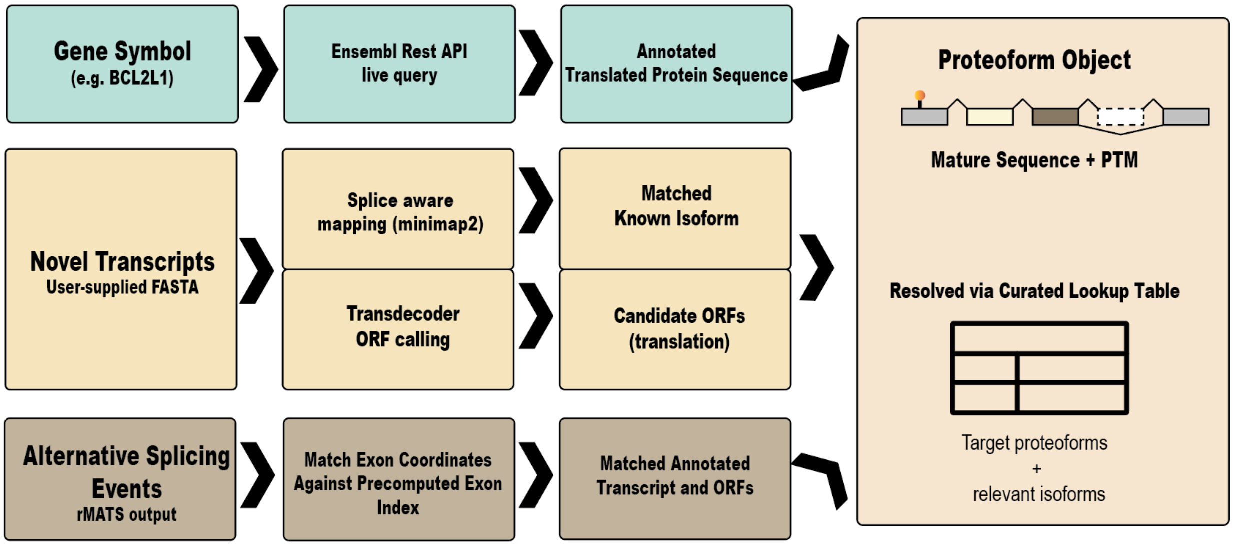


**Supplementary Fig. N1.** Detailed input-parsing and proteoform-inference workflow for ProteoformTracker's three input paths. Upper: (gene → isoform) resolves via a live, disk-cached Ensembl REST query. Middle: (novel transcript) runs minimap2 spliced alignment and TransDecoder ORF calling as two independent steps on the same input sequence. Lower: (rMATS event) matches each arm's exon coordinates against a precomputed exon-structure index to recover full-length annotated transcripts. All three converge on a shared PTM-annotation step before producing the common Proteoform object used by all downstream scoring.

### Supplementary Note 2: MS2 fragmentation-propensity modeling

**Data.** The fragmentation-propensity model was calibrated against two independent, publicly deposited HCD top-down proteomics datasets from MassIVE (MSV000094311 and MSV000098558), searched with TDPortal (Toby *et al.* 2019). From each dataset's confident proteoform-spectrum matches, every theoretical backbone cleavage position (b/y ion pair) of every matched proteoform was extracted and labeled *observed* (1) if a confidently matched fragment ion was reported at that position in the corresponding spectrum, or *observed* (0) otherwise, pooling to ≈2.7 million labeled bond-level observations across both datasets. For each bond, a fixed feature set was computed directly from sequence and PTM context: residue identity flanking the cleavage site (presence of proline immediately C-terminal, or aspartate immediately N-terminal, to the bond — both known HCD cleavage-enhancing motifs), distance to the nearer terminus (binned, capped at 6 residues), local basic-residue ("charge") density in a ±5-residue window around the bond, proximity to an annotated phosphorylation site, and the parent proteoform's total sequence length.

**Model.** Two scoring modes were fit from the same underlying features and labeled data, differing in whether proteoform length is included as a term.

*Calibrated (GLM) mode* is a single, jointly fit logistic regression of *observed* on the features above. The model outputs a fitted probability (0–1) directly via logistic regression. Refitting all terms in one joint model substantially improved cross-study generalization (see Results below) and is the version reported here. Proteoform length is included as a real, validated effect: under a fixed instrument duty cycle, longer proteoforms receive systematically sparser per-bond fragment-ion coverage, which the model captures via a clamped log-length term (clamped to the 5th–95th percentile of calibration-set proteoform lengths, 20–320 residues, to avoid extrapolating beyond what the data can support) — a proteoform-level effect applied identically to every bond in that proteoform, so it does not change the *relative* ranking of bonds within one proteoform, only the *absolute* score scale across proteoforms of different length.

*RF ranking mode* is a random forest fit on the same underlying pooled labels and sequence-context features (residue-pair, distance to the nearer terminus, local charge density, phosphorylation proximity) as the GLM, using each feature's raw continuous or binary form rather than the GLM's binned buckets, and deliberately excluding the length term. Because of the inherent deficiency of RF modes in extrapolation of protein lengths outside the range represented in these two datasets, the RF-based propensity score excludes protein length entirely and is instead intended for within-proteoform relative ranking.

**Results.** All GLM terms (residue-pair, terminal-position, charge-density, length, and phospho-proximity) were statistically significant in the final joint fit, with signs and approximate magnitudes replicating independently in each of the two datasets before pooling. Effect sizes for the two most consequential terms: proteoform length carried an odds ratio of 0.375 per doubling of length ($P < 2.2\times10^{-16}$), and phosphorylation proximity carried an odds ratio of 0.229 ($P < 2.2\times10^{-16}$), both indicating strong, real suppression effects, closely matching the effect sizes recovered by the earlier, separately-fit pieces this joint model replaces — the two approaches agree on the size of each individual effect; they disagree sharply on how well the effects predict held-out data once combined (see below).

Discrimination of the two scoring modes was evaluated the strict way: true cross-study holdout, refitting each model from scratch on one MassIVE dataset in its entirety and evaluating on the other, fully unseen dataset, in both directions (**Supplementary Fig. S7**), rather than resubstitution on the pooled data the models were fit on. Trained on MSV000094311 and evaluated on the independent MSV000098558 (baseline match rate 7.9%), calibrated GLM mode reached AUROC = 0.818 and AUPRC = 0.304, versus AUROC = 0.800 and AUPRC = 0.221 for RF ranking mode — GLM ahead on both metrics. In the reverse direction — trained on MSV000098558, evaluated on MSV000094311 (baseline match rate 5.6%) — the ranking flips on AUROC but not AUPRC: RF reached AUROC = 0.774 versus GLM's 0.755, while GLM retained the higher AUPRC (0.165 versus 0.145 for RF). Neither scoring mode dominates uniformly across both holdout directions, and both substantially and consistently exceed random-baseline discrimination (AUROC 0.5) throughout; RF's competitive-to-superior discrimination despite excluding the length term entirely indicates that the nonlinear interactions a tree ensemble can capture among the remaining sequence-context features (residue identity, unbucketed terminal distance, charge density, phospho-proximity) recover much of the separating information GLM's length term contributes — consistent with RF mode's intended role as a reliable ranking signal for long proteoforms whose absolute GLM scores the length penalty has suppressed.

Stringency-tier cutpoints for both scoring modes were selected by sweeping candidate thresholds against held-out fold-enrichment for matched fragment ions over the pooled dataset baseline match rate (6.3%); the deployed thresholds (GLM: Elevated > 0.10, High ≥ 0.20; RF: Elevated > 0.08, High ≥ 0.15) give approximately 3.1×/4.7× (GLM) and 2.5×/3.5× (RF) fold-enrichment over baseline. An initial three-tier scheme (adding a "Very High" cutpoint above the deployed High threshold in each mode) was evaluated but ultimately dropped for both scoring modes: at that stringency, few or no bonds cleared the threshold for the large majority of realistic-length proteoforms, making the tier practically unreachable rather than merely rare.
